# Development, field-testing and optimization of tools to quantify soil-transmitted helminths in fecal sludge from school pit latrines in Ethiopia

**DOI:** 10.64898/2026.02.11.705237

**Authors:** Abebaw Tiruneh, Zeleke Mekonnen, Sara Roose, Mio Ayana, Fiona Vande Velde, Emmanuel C. Mrimi, John Gilleard, Michael R. Templeton, Zewdie Birhanu, Jaco Verweij, Luc Coffeng, Bruno Levecke

## Abstract

**Background:** Surveys to monitor large-scale deworming programs against soil-transmitted helminthiases (STH) involve examination of stool samples from schoolchildren. These surveys are resource demanding and impact school activities. A potentially cost-saving alternative that does not involve children is to process fecal sludge samples from school pit latrines. To provide a proof-of-principle of latrine-based monitoring of STH programs, we optimized tools to collect fecal sludge and to quantify STH eggs in the samples.

**Methods:** First, we designed, developed and field-tested three locally made fecal sludge sampling prototypes. Second, we developed a modified egg-counting method and conducted spiking experiments to explore its analytical performance. Third, we estimated the variation in egg counts in fecal sludge samples collected from six primary schools in Ethiopia at different pit latrine depths used by boys and girls and by repeatedly examining samples. Finally, field data were used to inform an egg count simulation model to quantify this variation in egg counts and to determine the sampling and analysis strategies that resulted in surveys as precise as a stool-based survey.

**Results:** The modified fecal sludge sampling prototypes were generally successful, except for a few pit latrines with dried/solid sludge types and insufficient sludge volume. The egg-counting method had moderately high analytical sensitivity that varied across the consistency of the samples. The variation in egg counts was mainly explained by differences between squat holes followed by repeated fecal sludge sample processing. Latrine-based surveys were as precise as stool-based surveys only for *Ascaris* and when the intensity of infections was low.

**Conclusions:** We developed a sampling and diagnostic strategy that we will use in a follow-up study. This study will be conducted in Jimma Zone across 25 schools (52 children per school) and will compare the mean fecal sludge egg counts at school level with the STH prevalence in children.

**Author summary:** Progress of large-scale deworming programs are currently monitored through screening individual stool samples of schoolchildren. A latrine-based program monitoring, without active participation of schoolchildren and interruption of routine school activities, is a potentially cost-effective alternative. As a proof-of-concept, we developed a method to quantify worm eggs in fecal sludge samples, conducted spiking experiments to determine its analytical performance and applied the method across schools in Ethiopia to explore variation in egg counts (e.g., squat holes, depth of sample collection, and repeated analyses). Based on our findings, we then determined the sample collection and analysis strategy that results a latrine-based survey as precise as a survey based on screening individual stool samples. The analytical performance was moderately high but varied across the consistency of the samples. The variation in egg counts was mainly driven by variation between squat holes on the same pit latrine, and thus it is better to sample more squat holes (at least 3) than repeatedly processing the same fecal sludge sample (one examination is sufficient). We will now apply this sample collection and analysis across 25 schools (52 children per school) in Ethiopia and compare it with a survey based on screening individual stool samples.

## Introduction

Soil-transmitted helminthiases (STH) are caused by *Ascaris lumbricoides*, *Trichuris trichiura* and hookworms (*Necator americanus* and *Ancylostoma duodenale*). These diseases are transmitted through a sequence of events including (i) infected individuals excreting eggs laid by female adult worms into the environment through stool; (ii) the excreted eggs developing into infectious life stages; (iii) the life stages passively (oral uptake) or actively (skin penetration) entering individuals, and (iv) maturing to adult worms in the intestines. Because of this route of transmission, the diseases persist in settings where open defecation is frequently practiced, and where there is lack of access to clean water, basic sanitation, and low hygiene standards [1–3]. STH-attributable morbidities are mainly associated with moderate-to-heavy intensity (MHI) infections, resulting in malnutrition, physical and cognitive retardation in children, and anemia and negative birth outcomes (e.g., low birth weight) in women of reproductive age [2,4–6]. To reduce these morbidities, the World Health Organization (WHO) recommends large-scale deworming programs in endemic areas, during which anthelmintic drugs are periodically administered to at-risk populations [7,8], including but not limited to children.

Between 2010 and 2019, a significant reduction in the disease burden was observed (2.7 million DALYs in 2010 *vs.* 1.9 million DALYs in 2019) [9]. Encouraged by these global successes, WHO has now moved away from program coverage targets, and has defined new targets for 2030 that better reflect the maturity of the STH programs [10]. The targets include sustaining elimination of STH as a public health problem in endemic countries (prevalence of MHI infection in children <2% (**target #1**) and a 50% reduction of tablets of anthelmintic drugs needed in STH control programs (**target #2**). As both targets are dependent on the outcome of epidemiological surveys to verify an elimination status (target #1) or to guide decisions on scaling down or stopping the administration of tablets (target #2), close monitoring of these programs is of utmost importance [11].

Today, the recommended monitoring and evaluation (M&E) tool is based on screening of stool samples from 250 schoolchildren across five schools (50 children per school) within a district using Kato-Katz thick smear technique [2,10,11]. However, recent studies questioned whether this survey design allows for reliable program decision-making, and screening of individual stool samples remains resource intensive [12–15]. A substantial proportion of the cost of performing these surveys is related to the individual stool sampling (transportation and per diem), followed by the process to prepare and count eggs in a stool smear [16,17]. As such, major cost drivers of M&E surveys are the number of schools and children to be screened, the speed at which technicians can process a single sample, the number of samples that can be processed per day, and thus the number of sampling days [18,19]. Although pooling samples [17,20] has been considered a cost-saving strategy, this proved to be sobering. Indeed, a pooling strategy mainly reduces the laboratory time, and not the resources required to collect the individual stool samples [17]. Therefore, it is important to explore alternative surveillance tools to monitor STH control programs [21].

A potential cost-saving strategy that merits more research is monitoring the environmental contamination (e.g., soil, wastewater, and fecal sludge) rather than the infections in children. This could allow screening a larger sample of the population at the same operational cost, and ultimately in more reliable program decision-making [22–27]. Monitoring the environmental samples also takes away important operational obstacles that programs are currently facing (e.g., expedites ethical process for stool samples and avoids class interruptions during surveys). Yet to date, there is little to no evidence that monitoring the environment is a cost-efficient alternative to inform STH control programs [6].

In our previous work, we demonstrated that soil samples from school compounds (playground, around the latrines and classrooms), households and open markets were highly contaminated with worm life stages, including but not limited to those causing STH. The environmental contamination at school level was associated with the prevalence of any STH across random sample of the children at those schools [25]. Here too, we encountered some important operational challenges related to both sampling and diagnostic strategy. While our samples were easy to collect (a shovel was the main equipment), it is not clear what number of samples should be collected to allow for an accurate and precise assessment of the environmental contamination. Our diagnostic method had a moderate detection limit (50 eggs per 100 g of soil for *Ascaris* and *Trichuris* eggs using microscopy), but it required expensive equipment (price: 8,000.00 EUR). Based on these challenges, we moved away from sampling soil and want to explore the potential of examination of fecal sludge samples from pit latrines instead.

As a step to provide a proof-of-principle of a latrine-based M&E of STH control programs, this study aimed (i) to develop low-cost devices to collect fecal sludge samples, (ii) to develop a method to quantify STHs in fecal sludge samples under laboratory conditions, (iii) to field test both the sample collection devices and the egg-counting method, and (iv) to determine the sampling (the required number of sludge samples) and the analytical (number of preparations per sample and number egg counts per sample preparation) strategies to reliably assess the concentration of STH eggs in school pit latrines in Ethiopia.

## Methods

### Ethics statement

Ethical approval was obtained from the National Research Ethics Review Committee under The Ministry Education (REF NO: 8/143/732/25) based on a support letter from Institutional Review Board (IRB) of Jimma University, Ethiopia (Ref. No: JUIH/IRB/287/24). Similarly, ethical approval was received from Ghent University, Belgium (Ref. No: ONZ-2024-0364). Letters from Molecular Biology and NTD Research Center were distributed to each primary school involved in the study, and permission was sought from the schools’ administrations to collect fecal sludge samples from school pit latrines. We also obtained informed consent from household heads to collect fecal sludge samples from three household pit latrines for spiking experiment.

### Development of low-cost fecal sludge sampling devices

We performed an exploratory review of scientific literature and targeted Google searches to identify existing devices for *in situ* grab sampling of fecal sludge from pit latrines at different depths. Guided by two key criteria: (i) low production cost and (ii) feasibility of local manufacture using readily available materials, we assessed whether a suitable, readily available sampler could be used or whether a new device needed to be designed. Our review was non-systematic and intended to inform the selection or development of appropriate sludge sampling devices rather than to provide a comprehensive overview. Based on the insights gained, we designed prototype sampling devices. The suitability of the prototypes was subsequently evaluated under field conditions in seven primary schools (four schools in Jimma Town and one schools in Manna District, Southwest Ethiopia).

### Quantification of STH eggs from fecal sludge samples

To quantify STH eggs in fecal sludge samples, we adapted our previously developed and validated diagnostic method for soil samples [25]. Briefly, the method for soil samples was based on (i) mixing samples in a liquid phase containing detergents (tween 80), (ii) sieving samples over a tower of sieves that were automatically shaken and washed, and further concentrating the eggs through (iii) steps of centrifugation, filtration and flotation. For the fecal sludge samples, we simplified the procedure by replacing the automated sieving and washing machine with the ‘Fluke Catcher’ and using only water instead of detergent solutions.

The Fluke Catcher is a hand-held stack of three sieves (sieve 1: 189 μm, sieve 2: 104 μm and sieve 3: 59 μm; all with diameter of 6 cm, and a total height of 32 cm height). It is made commercially available by Provinos (price: 109.50 EUR) [28] to detect *Fasciola* eggs in feces of ruminants. Because STH eggs pass through all sieves of the Fluke Catcher, we decided to use an additional metal sieve of 10 cm (diameter) x 4 cm (height) and a mesh size of 20 μm (price: 369 EUR) [29]. **Fig 1** briefly describes the procedures, and a detailed standard operating procedures (SOP) is available in **Info S1.**

**Fig 1.**
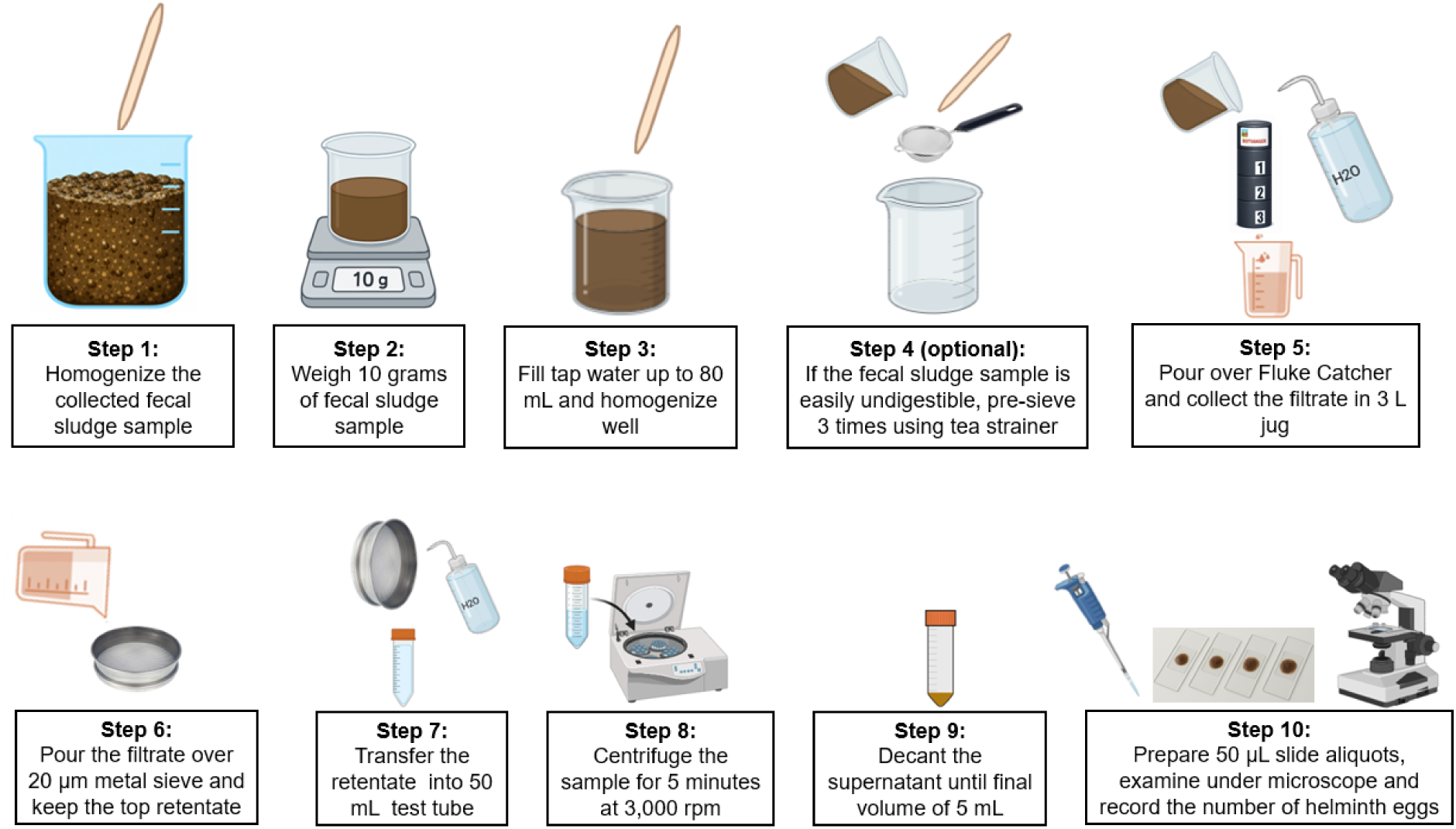
An overview of the procedure to recover and quantify STH eggs in fecal sludge samples. The procedure of consists of 10 consecutive, including: homogenizing the collected fecal sludge sample in the original container (fecal sludge sample bucket) to uniformly distribute of eggs in the fecal sludge **(Step 1)**, weighing 10 grams of fecal sludge sample using digital balance in 100 mL beaker **(Step 2)**, adding tap water until 80 mL and mixing the suspension **(Step 3),** straining the sample 3 times using tea strainer to withhold the large debris **(Step 4)**, straining the sample over Fluke Catcher and collecting the filtrate in a 3 L jug **(Step 5)**, straining the filtrate over 20 µm metal sieve **(Step 6)**, transfer the retentate into a 50 mL test tube including the rinsing water of the metal sieve **(Step 7)**, centrifuging the test tube for 5 minutes at 3,000 rpm **(Step 8)**, decanting the supernatant up to a final volume of 5 mL **(Step 9)** and transferring 50 µL onto a slide and counting the STH eggs using a compound microscope (10×10 magnification) **(Step 10)**.

### Spiking experiments

For the spiking experiments, we collected two fecal sludge samples from pit latrines of three households and screened them using our modified Fluke Catcher method. To ensure that these fecal sludge samples were negative for helminth eggs, each sample was processed four times and three slides (each representing a volume of 50 µL) were examined for each sample preparation. To simulate the three types of fecal sludge consistencies, we measured the total solids (TS) of the collected two fecal sludge samples and diluted them (when needed) with tap water to obtain semi-solid (TS: 15.1-25%), slurry (TS: 5.0-15.0%) and liquid (TS: <5%) samples [30]. The *Ascaris* eggs were obtained from both *Ascaris*-positive stool and fecal sludge samples using our soil-straining flotation method and stored in a stock volume of 3 – 5 mL at 2 - 8 °C. We then examined the 50 µL of the stock solution in triplicate to estimate the concentration (number of purified *Ascaris* eggs per 1 µL).

To evaluate the analytical performance of our modified Fluke Catcher method, we spiked 50, 100, 200 and 400 purified *Ascaris* eggs into negative fecal sludge samples of different types; semi-solid, slurry and liquid. The spiking was repeated eight times for each combination of number of spiked *Ascaris* eggs and sludge type, resulting in 96 spiked samples (8 replicates x 4 number of spiked eggs x 3 types of sludge) and 384 worm egg counts (96 spiked samples x 4 slides of 50 µL). All samples were then processed following the SOP for the modified Fluke Catcher to quantify STH eggs in sludge sample as described above. To avoid any systematic error, the spiking experiment was randomized for number of *Ascaris* eggs spiked and type of sludge. The laboratory staff who performed the sample processing and microscopy examinations were blinded.

### Field evaluation of the fecal sludge sampling devices and the modified Fluke Catcher method

#### School selection

We collected fecal sludge samples from six primary schools in Jimma Town, Southwest Ethiopia. The schools were selected based on their involvement in previous epidemiological, drug efficacy trial and diagnostic performance assessment surveys [20,25,31,32]. For each school, we first identified and inspected all pit latrines available in the school compound. When sampling fecal sludge was not possible from at least one pit latrine, for example when the sludge had dried out, we proceed to collect samples from the next school.

#### Fecal sludge sampling and processing

For pit latrines shared by both boys and girls, we collected fecal sludge samples from two latrine squat holes per sex. When pit latrines were separated for boys and girls, we randomly sampled two latrine squat holes from each (**Fig 2**). Whenever possible, one superficial (depth up to 0.2 m) and one deep (depth >0.5 m) fecal sludge samples [33] were collected for each latrine squat hole following the SOPs for sludge sample collection and analysis **(Info S1)**.

**Fig 2.**
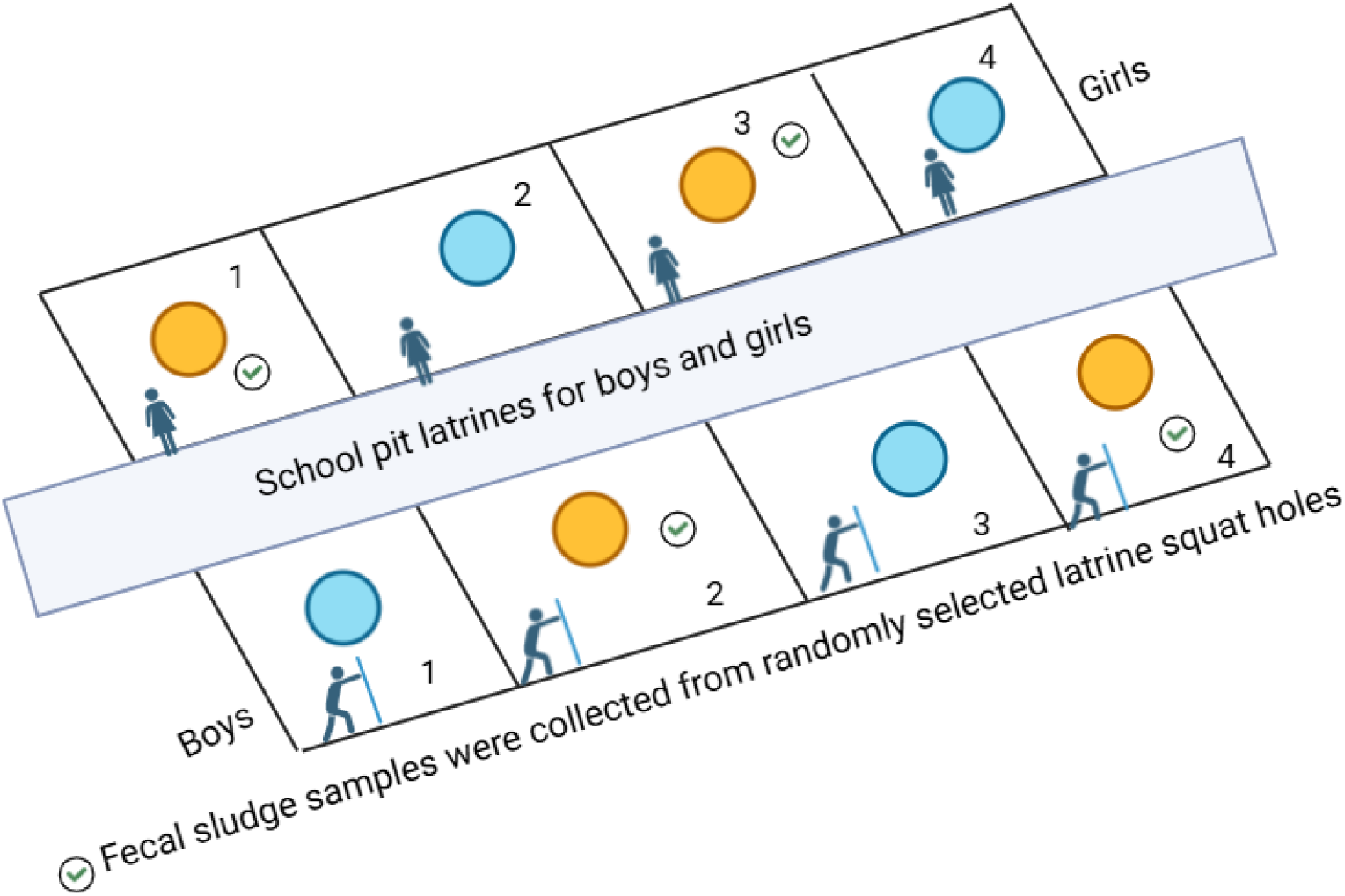
The selection of squat holes for the collection of fecal sludge samples. All pit latrines in each school compound were identified, and their eligibility was checked (eligible implied that pit latrines were used and fecal sludge sample collection was feasible). Eligible pit latrines were sketched on paper and assigned a numbers (1 to the total number of squat holes). Two holes for each sex (boys and girls) were randomly selected (yellow circles with a tick mark) and fecal sludge samples were collected.

The volume of the collected fecal sludge samples ranged from 0.5 L to 1.3 L. Environmental and personal safety were maintained at all steps of the fecal sludge sample collection procedures. A total of 40 (24 superficial and 16 deep) fecal sludge samples were collected and transported to Jimma University Molecular Biology and NTD Research Center using a cooler box. Once arrived at the laboratory, each sample was repeatedly processed (up to four sample preps per sample) as described in previous sections, including the assessment of the TS. All results were expressed as egg counts per gram of TS (EPG_TS_) for each of the STH separately.

### Statistical analysis

#### Analytical performance of the modified Fluke Catcher method

We evaluated the performance of the modified Fluke Catcher method based on (i) the analytical sensitivity (the number of spiked *Ascaris* eggs that resulted in a positive test result with a probability of at least 95%) and (ii) the egg recovery rate. For both parameters, we used a regression model for count data including a random intercept for processed fecal sludge samples to account for correlation of counts within the sets of four repeated slides. To let coefficients represent the (logarithm of the) relative difference in expected and observed counts (i.e., the egg recovery rate), we included an offset for the natural logarithm of the expected number of eggs in a slide. As fixed effects, we considered the sludge type (liquid, slurry, or semi-solid) and the number of spiked eggs (50, 100, 200, or 400) as categorical predictors. The final selection of fixed effects was based on the Akaike Information Criterion (AIC), where we considered an improvement in AIC as indicative of better model performance. Based on this model, for each sludge type and a range of number of spiked eggs, we calculated the analytical sensitivity in terms of the probability of detecting at least 1 egg (averaged over all the estimated random intercepts). The model was implemented in R (version 4.5.1), using the package *glmmTMB* (version 1.1.13). The script for the analysis is provided at https://gitlab.com/luccoffeng/echolatrine.

#### Prevalence and abundance of STH life stages in field samples

We graphically explored the prevalence (%) and abundance (EPG_TS_) of STH life stages in fecal sludge samples across schools, depth of sampling (deep *vs.* superficial), and usage (boys *vs.* girls). For this, we determined (i) the prevalence (proportion of samples that test positive expressed in %) and the corresponding 95% confidence interval (CI) based on the binomial exact test and (ii) the median and interquartile range of the concentration (expressed in EPG_TS_) for each of the three factors.

#### Sources of variation in egg counts across field samples

To assess and quantify the contribution of different sources (e.g., schools, latrine squat holes, sample preps, repeated slides, and microscopists) to the variation in raw slide level egg counts in fecal sludge samples, we adapted an existing model for variance decomposition analysis of *S. mansoni* egg counts in humans [34]. With this model, we decomposed the variability of egg counts across sources using a multi-level Poisson regression model for all three species combined, using fixed effects to capture differences between schools, species, sex-specific latrine squat holes, and sampling depth. Fecal sludge type was not included as a predictor as it was always the same within five of the six schools. To correct for variation in absolute egg counts due to variation in consistency of faecal sludge samples, we included an offset for TS of each sample processed; consequently, all fixed effects represent the (logarithm) of the relative difference between samples in terms of EPG_TS_. We used gamma-distributed random effects to capture variation in egg counts between pit latrine squat holes, repeated sample preps and microscopists. In a sensitivity analysis, instead of gamma distributions we used the lognormal distribution for random effects. The contribution of pit latrine squat holes, repeated sample preps and microscopists (i.e., the random effects) to the overdispersion of egg counts within schools was expressed in terms of the coefficient of variation or *CV* of each random effect (see **Info S3** for technical details).

The regression model was implemented in a Bayesian framework, using the probabilistic programming language Stan (cmdstan version 2.35.0) and the R package *cmdstanr* (version0.9.0.9). Model parameters were sampled using 4 Markov chains, each with 1,000 warm-up and 1,000 sampling iterations, such that the effective number of samples from the posterior was at least 500 for all parameters. A description of the regression model in terms of equations is provided in **Info S2**; the corresponding R code to perform the analysis is provided at https://gitlab.com/luccoffeng/echolatrine.

#### Determination of the sampling strategy to reliably assess the concentration of STH eggs in pit-latrines

In this study we did not measure infection in children and could therefore not compare the reliability of school-level estimates based on testing of fecal sludge *vs.* stool samples from children. Instead, we defined the reliability of latrine-based estimates of infection levels in terms of the CV of the estimated school-level mean EPG_TS_ and compared this to the expected CV of stool results from children, based on a simulation exercise. Here, we assumed that, like in our field study, the sample collection from latrines would not be affected by heavy dilution from for instance rainwater or flooding.

Egg counts in fecal sludge samples were simulated using the framework described above, quantified based on the spiking experiment and field data. Egg counts in stool samples from children were simulated using a similar and previously published simulation framework to determine the most cost-efficient survey design to make accurate program decisions [35]. In the present study, we assumed that 52 children would be sampled and tested, and this is based on the results of Kazienga A., *et al*. [35], indicating that this is the sample size that allows for reliable decision making. Simulated survey results for latrines and children were both summarized in terms of the expected relative variability (expressed in CV) of the estimated school-level mean EPGs. These school-level CVs were simulated and calculated for a range of assumed true infection levels in the school. By necessity, we assumed that true EPG_TS_ in fecal sludge is the same as the true mean EPG in stool from children. As for the observed egg counts in simulated samples, we assumed that the egg recovery in stools from children was 100% and 50% in latrine samples, where the latter was based on the estimated egg recovery rate of the Fluke Catcher method in the egg spiking experiment. These assumptions will need to be revisited in future studies that sample both children and latrines from the same schools. The code to conduct the simulation experiment can be found at https://gitlab.com/luccoffeng/echolatrine.

## Results

### Field evaluation of fecal sludge sampling devices

The exploratory search of the scientific literature and targeted Google searches identified a range of self-made, commercially available, and modified commercial devices for fecal sludge sampling [36–44]. Based on the insights gained from this search, we identified two key criteria: (i) low production cost and (ii) feasibility of local manufacture using readily available materials. Based on this, we designed three prototype devices for *in situ* faecal sludge grab sampling (**Fig 3**). The design accounted for variation in pit latrine squat hole sizes, pit depths and fecal sludge types (liquid, slurry and semi-solid). Each device had a retractable stick with a maximum length of 5 m that allowed sampling at required depth and a container with a capacity of 1 L to 2 L was attached to one end of the stick. Two devices consist of an open sample container with diameters of 9 cm (**Fig 3A**) and 10 cm (**Fig 3B**). The third device used a closed container with a diameter of 10 cm, and its lid could be opened or closed by pulling a string (**Fig 3C**) to collect samples at a specific depth within a pit latrine.

**Fig 3.**
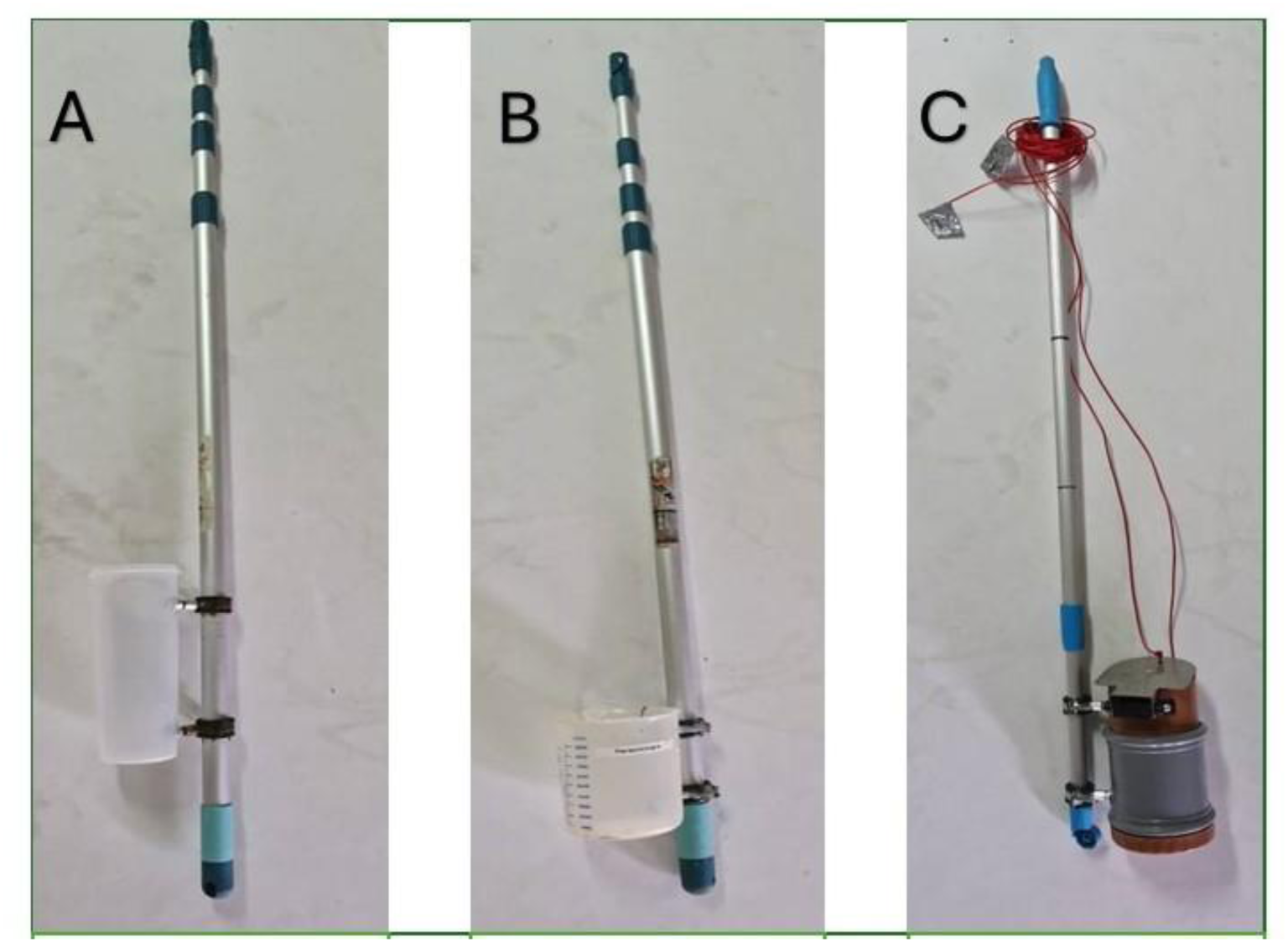
In-house fecal sludge sample collection devices. **Panels A** and **B**, represent devices with an open sample container with different diameters (Panel A: 9 cm and Panel B: 10 cm). **Panel C** represents a sampling collection device that that could be opened/closed by pulling a string (C). The diameter of the container equals 10 cm.

During field evaluation, the following main operational challenges were identified: narrow squat holes, the presence of foreign objects (e.g., plastic waste) within pit latrines, dried or solid fecal sludge, and sludge at greater depths. Based on these field observations, the prototypes were modified for use in the remainder of the study. To accommodate narrow squat holes, a narrower sampling container was secured with plastic cable ties. To enable sampling from deep pit latrines (>4 m), an additional extension stick was added and secured with duct tape. Despite these modifications, the devices remained unsuitable for sampling dried or solid fecal sludge and for latrines with insufficient sludge volume.

### Analytical performance of the modified Fluke Catcher method

The TS of the two collected fecal sludge samples were 21.5% and 17.9%. Subsamples of these fecal sludge samples were diluted with tap water to generate three slurry (TS = 7.7%, 9.7% and 10%) and three liquid (TS = 2.6%, 3% and 4.8%) fecal sludge samples. In total eight samples were spiked and analyzed using our modified Fluke Catcher method. Variability of egg counts in repeated slides closely followed a Poisson distribution (**S1 Fig)**. A numeric summary of the experiment is provided in **Table S1**.

Overall, the recovery rate of *Ascaris* eggs by our modified Fluke Catcher method was significantly higher in liquid sludge (85%, 95% CI: 74%–98%) than in slurry (65%, 95% CI: 55%–76%) and semi-solid sludge (56%, 95% CI: 47%–67%) (**Fig 4**). The egg recovery rate did not differ significantly between the slurry and semi-solid sludge. The number of spiked eggs was not a significant predictor of the egg recovery rate, either on its own (AIC = 1,068.4), or in combination with sludge type (AIC = 1,058.6), or as an interaction with sludge type (AIC = 1,067.0).

**Fig 4.**
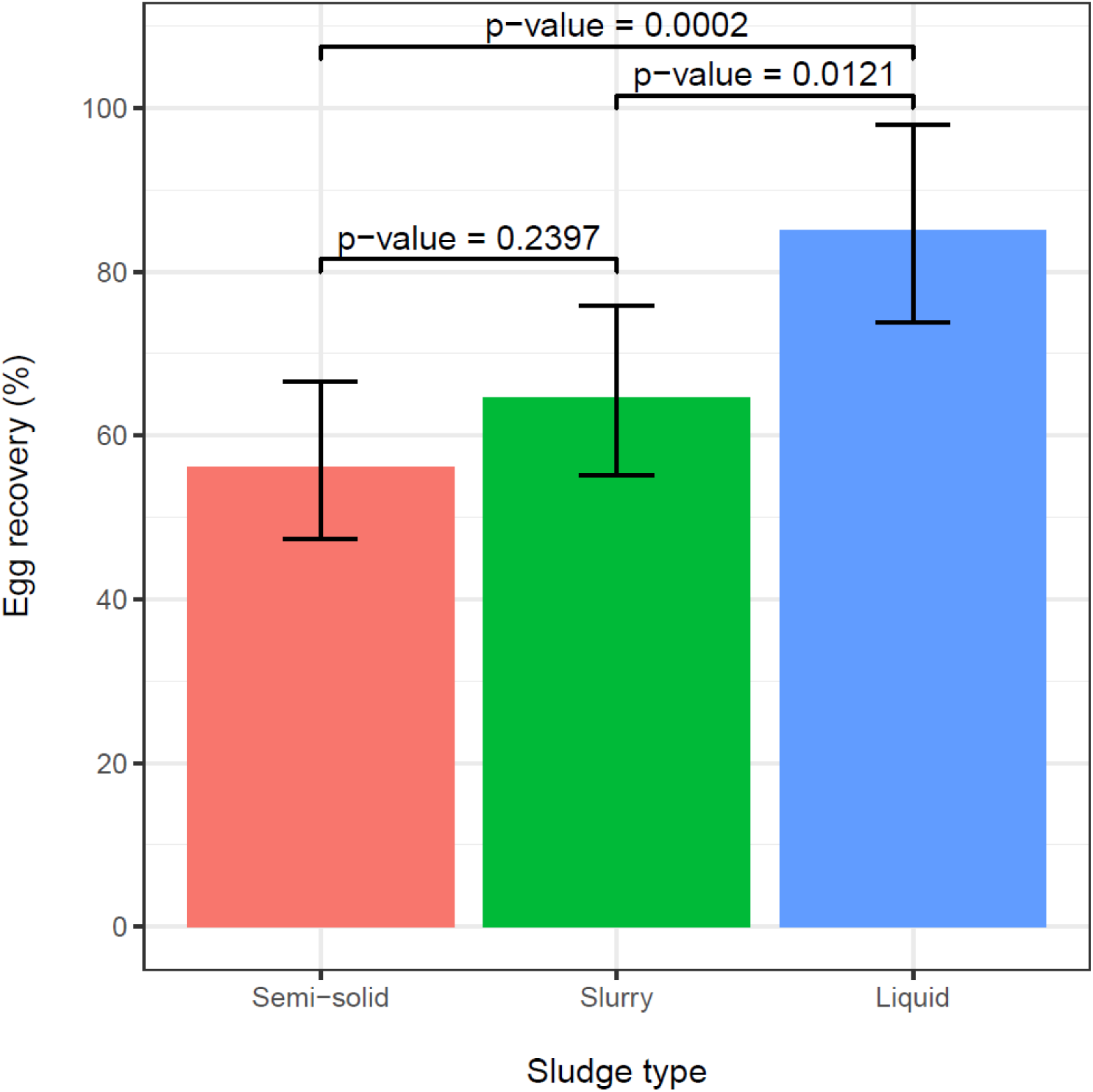
Estimated egg recovery rate from fecal sludge spiked with known quantities of *Ascaris* eggs. Each bar represents the average egg recovery rate across the sludge samples that were spiked with 50, 100, 200 or 400 eggs. Error bars indicate the 95% confidence interval around the point estimates. Results are based on a Poisson regression model with a random intercept per sludge sample (from which four repeated slides were taken), an offset for the natural logarithm of the expected number of eggs, and a fixed effect for sludge type.

Based on the estimated egg recovery rate, we calculated the analytical sensitivity of the Fluke Catcher method in terms of the probability of finding at least one egg (**Fig 5**). Generally, when four slides (each 50 µL) were examined per sample, the probability of finding at least one egg was ≥95% (dashed horizontal black line in **Fig 5**) when ≥ 90 eggs were spiked for liquid samples, ≥116 eggs for slurry samples, and ≥135 spiked eggs for semi-solid samples (dashed vertical lines). When reducing the number of slides, the analytical sensitivity dropped for all three fecal sludge samples.

**Fig 5.**
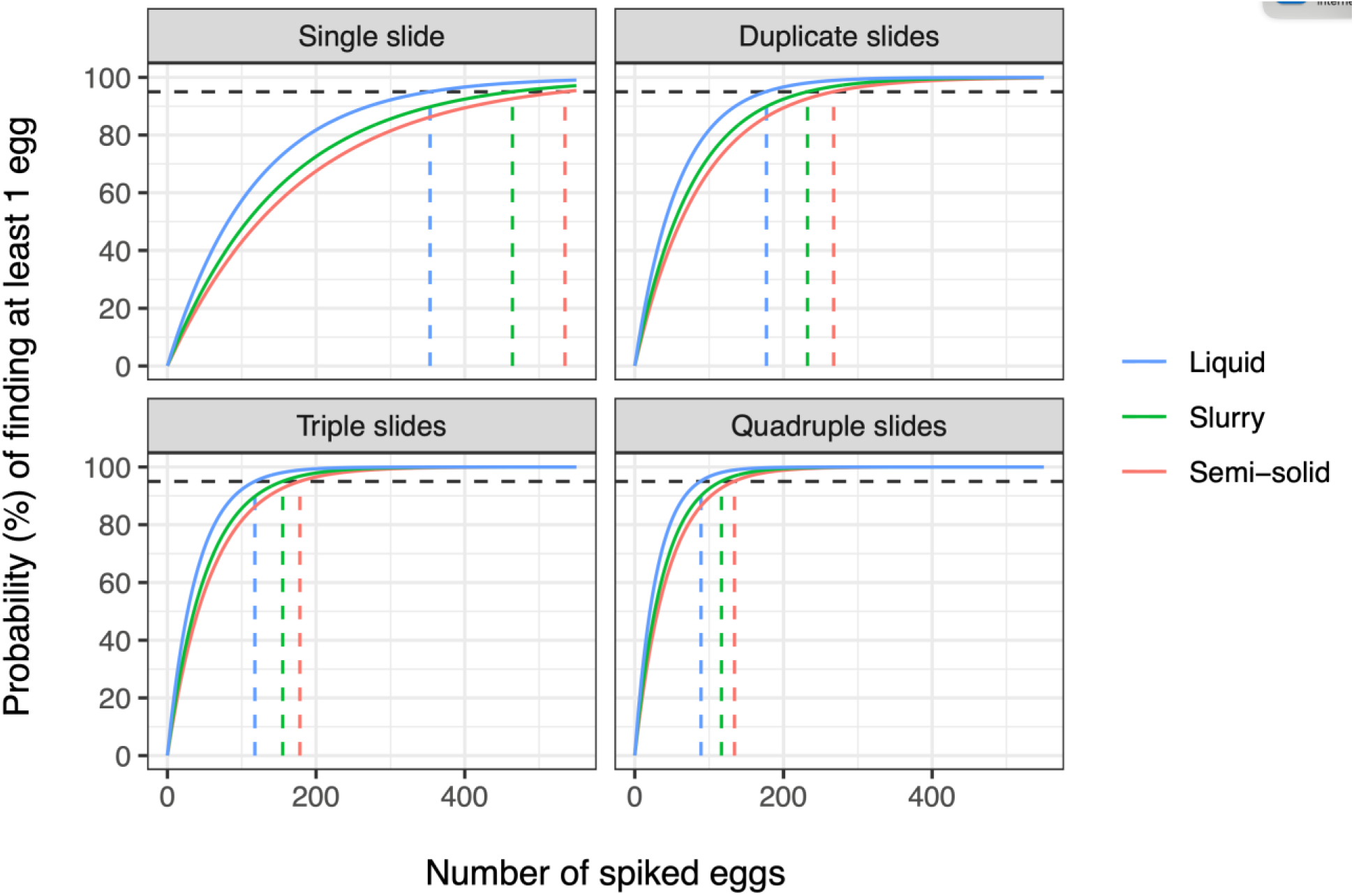
The analytical sensitivity of the modified Fluke Catcher methods to detect *Ascaris* eggs in fecal sludge samples. This represents the predicted probability of finding at least one egg for a wide range of spiked eggs for three fecal sludge types and different numbers of slides (each of 50 µL).

### Prevalence and abundance of STH life stages in fecal sludge samples

A total of 352 slides (each 50 µL) were prepared from 88 sample preps of 10 grams of fecal sludge from 40 samples collected at varying depths in pit latrines across six schools (**Table 1**). The consistency of fecal sludge samples varied across latrines and holes, with most samples being of the semi-solid type (n = 28) and the remainder being either slurry (n = 4) or liquid (n = 8). Superficial and deep fecal sludge samples from the same pit latrine squat hole were always of the same consistency type.

**Table 1.** Overview of collection and processing of fecal sludge samples from 8 pit latrines across 6 schools.

| School | Number of sampled pit latrines | Pit latrine type | Total number of holes sampled | Sampling depth | Depth of deep samples (cm from sludge surface) | Repeated number of sample preps of 10 g | Number of repeated slides | Total number of slides |
| --- | --- | --- | --- | --- | --- | --- | --- | --- |
| Dilfire | 1 | Common | 4 | Superficial + deep | 30–65 | 1 or 4* | 4 | 16+64* |
| Ginjo | 2 | Sex-specific | 4 | Superficial | - | 1 | 4 | 16 |
| Hirmata | 2 | Sex-specific | 4 | Superficial + deep | 30–85 | 4 | 4 | 128 |
| Jimma | 1 | Common | 4 | Superficial | - | 4 | 4 | 64 |
| Kito | 1 | Common | 4 | Superficial + deep | 70–100 | 1 | 4 | 32 |
| Mendera | 1 | Common | 4 | Superficial + deep | 50 | 1 | 4 | 32 |
| **Total** | **8** | **-** | **24** | **-** | **30–100** | **-** | **-** | **352** |

\* For two of the four holes (one for each sex), only a single sample prep of 10 g of sludge was processed per sampling depth, leading to 16 slides (2 holes x 2 depths x 1 sample prep x 4 slides); for the other two holes (one per sex), four repeated sample preps of 10 gr of sludge were processed per sampling depth, leading to 64 slides (2 holes x 2 depths x 4 sample preps x 4 slides).

Across the 88 fecal sludge sample preps (352 slides), we observed several helminth life stages of medical importance, including *Ascaris, Trichuris*, *Schistosoma mansoni* and others such as *Hymenolepis nana*, *Enterobius vermicularis,* and *Taenia* spp. We also observed hookworm-like eggs and unknown larvae but did not use any molecular tools to further identify these. *Ascaris* eggs were the most prevalent (343 / 352 slides; 97%) and abundant (6,240 mean EPG_TS_, including egg-negative slides). *Trichuris* was present in just over half of the slides (205 / 352; 58%) with lower abundance (133 mean EPG_TS_). *S. mansoni* eggs were the least prevalent (48 / 352; 14%) and abundant (17 mean EPG_TS_). Prevalence of *Ascaris* was 100% across five of the schools’ latrines, and 72% in one school latrine (Kito; **Fig 6A**). Prevalence of *Trichuris* eggs in slides varied considerably across schools’ latrines (ranged 3%–80%), as did the prevalence of *S. mansoni* eggs (0%-91%). For all three helminth species, abundance of eggs varied considerably across schools’ latrines, even when the prevalence was 100%, as for *Ascaris* (**Fig 6D**). However, for all three species, the prevalence and abundance of eggs did not seem to vary much between boys’ and girls’ latrines (**Figs 6B** and **6E**) or superficial and deep fecal sludge samples (**Figs 6C** and **6F**).

**Fig 6.**
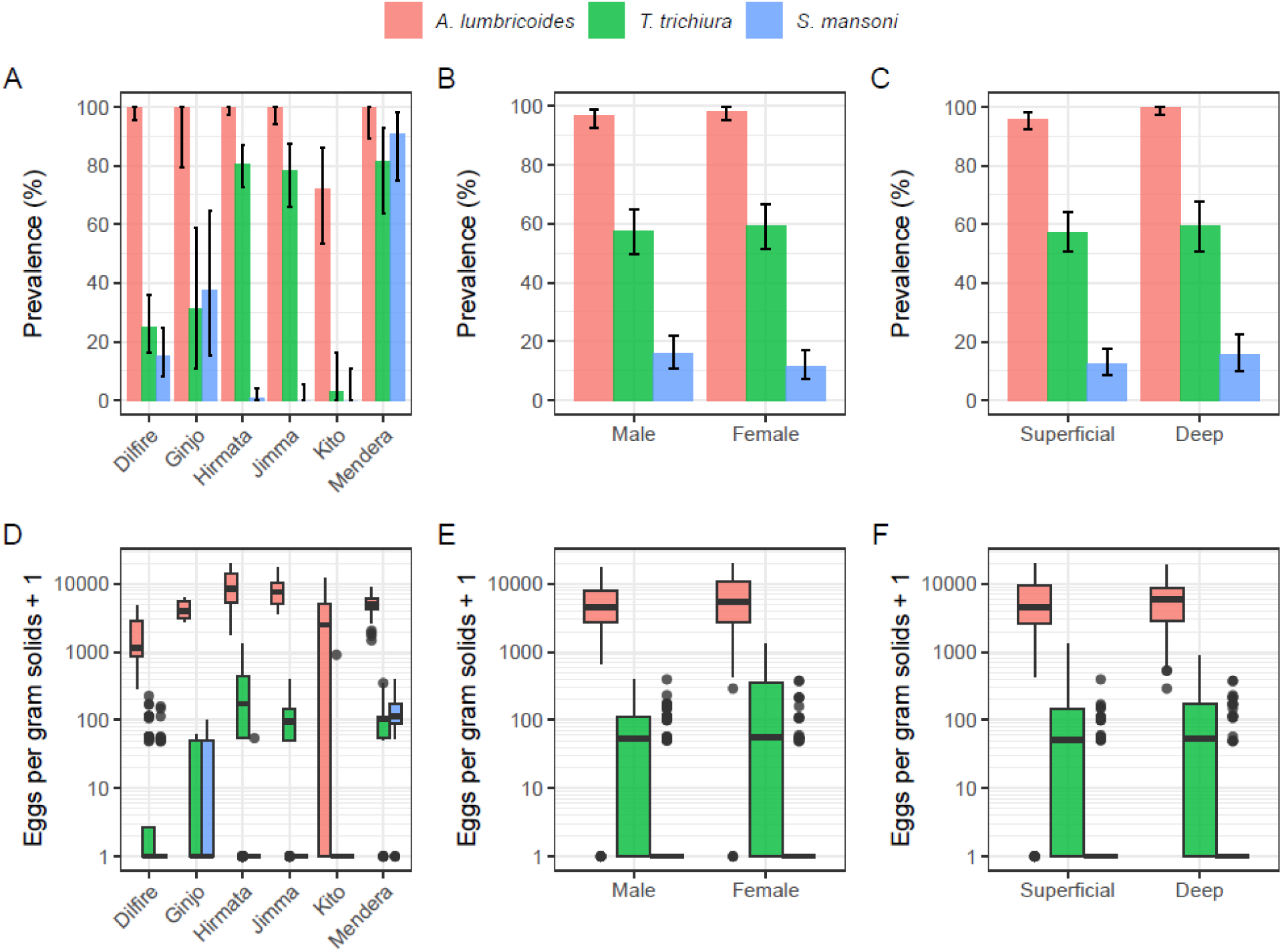
Prevalence and abundance of helminth eggs across 352 slides from six schools. The top row represents the prevalence (percentage of slides) of *A. lumbricoides*, *T. trichiura* and *S. mansoni* eggs across 6 schools (**Panel A**), sex-specific latrines (**Panel B**), and the depth at which samples were collected (**Panel C**). The boxplots in the bottom row represent the abundance of egg (eggs per gram solids detected in 50 µL of sample prep) across schools (**Panel D**), sex-specific latrines (**Panel E**), and the depth at which samples are collected (**Panel F**). Boxplots represent the median (thick horizontal line) and the first and third quartiles (lower and upper bound of each box). The whiskers (thin vertical lines) extend from the first or third quartile to the lowest or largest value, respectively, no further than 1.5 times the interquartile range. Data beyond the end of the whiskers are plotted individually (black bullets). Note that the y-axes of panels D-F are logarithmic and represent the egg abundance plus one (to include zero values in the plot). Further, note that some of the boxplots for *S. mansoni* do not show blue because the median and first and third quartiles are all the same (horizontal thick black line segments) due to the high number of zero egg counts.

### Sources of variation in helminth egg counts across fecal sludge samples

To inform the structure of the count regression model, we visually assessed the level of overdispersion in slide level egg counts (variance relative to the mean). Overdispersion of egg counts was the strongest across pit latrine holes from the same school, with the variance across holes consistently exceeding the school-level mean egg count (**Fig S2**). In contrast, the variance in egg counts was very similar to the mean count across slides and across repeated sample preps (**Fig S2**). The only exception was *Ascaris*, for which the variance tended to exceed the mean count for the higher egg counts. A visual inspection of the raw slide-level count data () suggested that intra-operator variability might have contributed to overdispersion in slide-level egg counts for *Ascaris*. Out of each set of four repeated slides, the two highest and two lowest counts were often performed by different microscopists (i.e., non-overlapping counts). Of the 88 sets of four repeated slides, 80 sets were read by two microscopists. Within these 80 sets, the frequency of non-overlapping counts by the different microscopists was significantly higher than the expected 33% (1/3): 54% for *Ascaris* (binomial exact *p*-value <0.001), indicating that the presence of more than one microscopist significantly contributed to variation in egg counts. A similar pattern was found for *Trichuris* (64%, *p*-value <0.001) and *S. mansoni* (86%, *p*-value <0.001). Based on these observed patterns in variability of egg counts, we specified a multi-level count regression model with random effects capturing variation between holes, samples, slides, and microscopists, as described in the methods section.

As only 5 microscopists were involved in the counting of eggs, and none of them seemed to systematically over- or underestimate counts across the sets of 4 slides (each 50 µL), the random effect for microscopist was assumed to be independent across different sets of slides.

For all three parasite species, the regression model indicated that egg counts did not vary notably between pit latrine holes for boys and girls: the relative differences in parasite-specific EPG_TS_ in pit latrines for females vs. males (reference) were 1.1 (*Ascaris*; 95% Bayesian credible interval (BCI): 0.68–1.55), 0.87 (*Trichuris*; 0.5–1.4), and 0.7 (*S. mansoni*; 0.3–1.3). Sex was therefore not retained in the model as a predictor. Sampling depth was kept in the final model, as *Ascaris* eggs were 42% more abundant in deep fecal sludge samples than in superficial samples (95% BCI: 21%–66%). *Trichuris* and *S. mansoni* eggs also tended to be more abundant at deeper sampling depths, although the 95% BCI of these differences did span 0% (*Trichuris*: 11%, 95% BCI: −13%–38%; *S. mansoni*: 63%, 95% BCI: −3%–150%). Across all three helminth species together, deep fecal sludge samples contained 34% more eggs (95% BCI 18%–53%) than superficial samples.

The model further confirmed that within schools’ latrines and per species, the overall level of overdispersion in egg counts (*CV*_*total*_ = 0.72) was mostly driven by variation between pit latrine squat holes (*CV*_*J*_ = 0.56). Variation between sample preps of 10 g was the second most important source of variation (*CV*_*S*_ = 0.33) and variation between microscopists was the least important (*CV*_*M*_ = 0.18). **Table 2** summarizes these findings. This pattern held in a sensitivity analysis using a multi-level Poisson model with log-normal-distributed random effects (*CV*_*total*_ = 0.81, *CV*_*J*_ = 0.65, *CV*_*S*_ = 0.34, *CV*_*M*_ = 0.18).

**Table 2.** Model-estimated parameter values for variance decomposition of egg counts in fecal sludge samples from pit latrines.

| Parameter | Gamma-distributed random effects |  |  | Log-normal-distributed random effects |  |  |
| --- | --- | --- | --- | --- | --- | --- |
|  | Posterior mean | 95% Bayesian credible interval |  | Posterior mean | 95% Bayesian credible interval |  |
| <u>Mean EPG<sub>TS</sub> for *A. lumbricoides*\*</u> |  |  |  |  |  |  |
| Dilfire | 2,501 | 1,439 | 4,142 | 1,943 | 1,136 | 3,061 |
| Ginjo | 5,179 | 2,975 | 8,577 | 4,519 | 2,479 | 7,304 |
| Hirmata | 8,418 | 5,207 | 13,126 | 6,926 | 4,065 | 10,751 |
| Jimma | 8,661 | 5,336 | 13,664 | 7,665 | 4,630 | 11,875 |
| Kito | 3,111 | 1,773 | 5,064 | 2,489 | 1,374 | 3,985 |
| Mendera | 4,727 | 2,780 | 7,547 | 4,204 | 2,421 | 6,707 |
| <u>Mean EPG<sub>TS</sub> for *T. trichiura*\*</u> |  |  |  |  |  |  |
| Dilfire | 26 | 14 | 45 | 18 | 9 | 32 |
| Ginjo | 31 | 13 | 62 | 26 | 11 | 50 |
| Hirmata | 254 | 153 | 401 | 158 | 92 | 249 |
| Jimma | 110 | 65 | 174 | 92 | 53 | 147 |
| Kito | 60 | 17 | 136 | 53 | 15 | 115 |
| Mendera | 114 | 63 | 190 | 97 | 53 | 159 |
| <u>Mean EPG<sub>TS</sub> for *S. mansoni*\*</u> |  |  |  |  |  |  |
| Dilfire | 12 | 6 | 21 | 10 | 5 | 18 |
| Ginjo | 29 | 12 | 53 | 24 | 10 | 46 |
| Mendera | 107 | 58 | 182 | 93 | 49 | 151 |
| <u>Relative difference between deep versus superficial samples</u> |  |  |  |  |  |  |
| *A. lumbricoides* | 1.42 | 1.21 | 1.66 | 1.43 | 1.22 | 1.65 |
| *T. trichiura* | 1.11 | 0.87 | 1.38 | 1.11 | 0.86 | 1.39 |
| *S. mansoni* | 1.63 | 0.97 | 2.50 | 1.62 | 0.97 | 2.48 |
| <u>Coefficient of variation (CV)</u> |  |  |  |  |  |  |
| Total CV | 0.72 | 0.61 | 0.85 | 0.81 | 0.66 | 1.00 |
| CV for holes | 0.56 | 0.45 | 0.70 | 0.65 | 0.49 | 0.85 |
| CV for samples | 0.33 | 0.27 | 0.39 | 0.34 | 0.28 | 0.42 |
| CV for microscopists | 0.18 | 0.15 | 0.21 | 0.18 | 0.15 | 0.22 |

\*Mean egg count per gram total solids in superficial fecal sludge samples. Estimates from the gamma-distributed random effects model represent arithmetic means, whereas values for the log-normal-distributed random effects model represent geometric means. Average egg counts for *S. mansoni* were only estimated for three schools (Dilfire, Ginjo, Mendera) as no eggs were detected in other schools (Jimma, Kito) or only in a single sample (Hirmata).

### Fecal sludge sampling strategy to reliably assess the concentration of STH eggs in pit latrines

We determined an optimized fecal sludge sampling strategy based on the association between the variability in measured school-level average egg intensity (expressed as CV) and the true school-level mean egg density (a proxy of endemicity) for different scenarios of diagnostic effort (number of sample preps) and sampling effort (number of squat holes sampled). We are assuming one slide per sample prep. **Fig 7** illustrates two aspects of this association (black lines).

**Fig 7.**
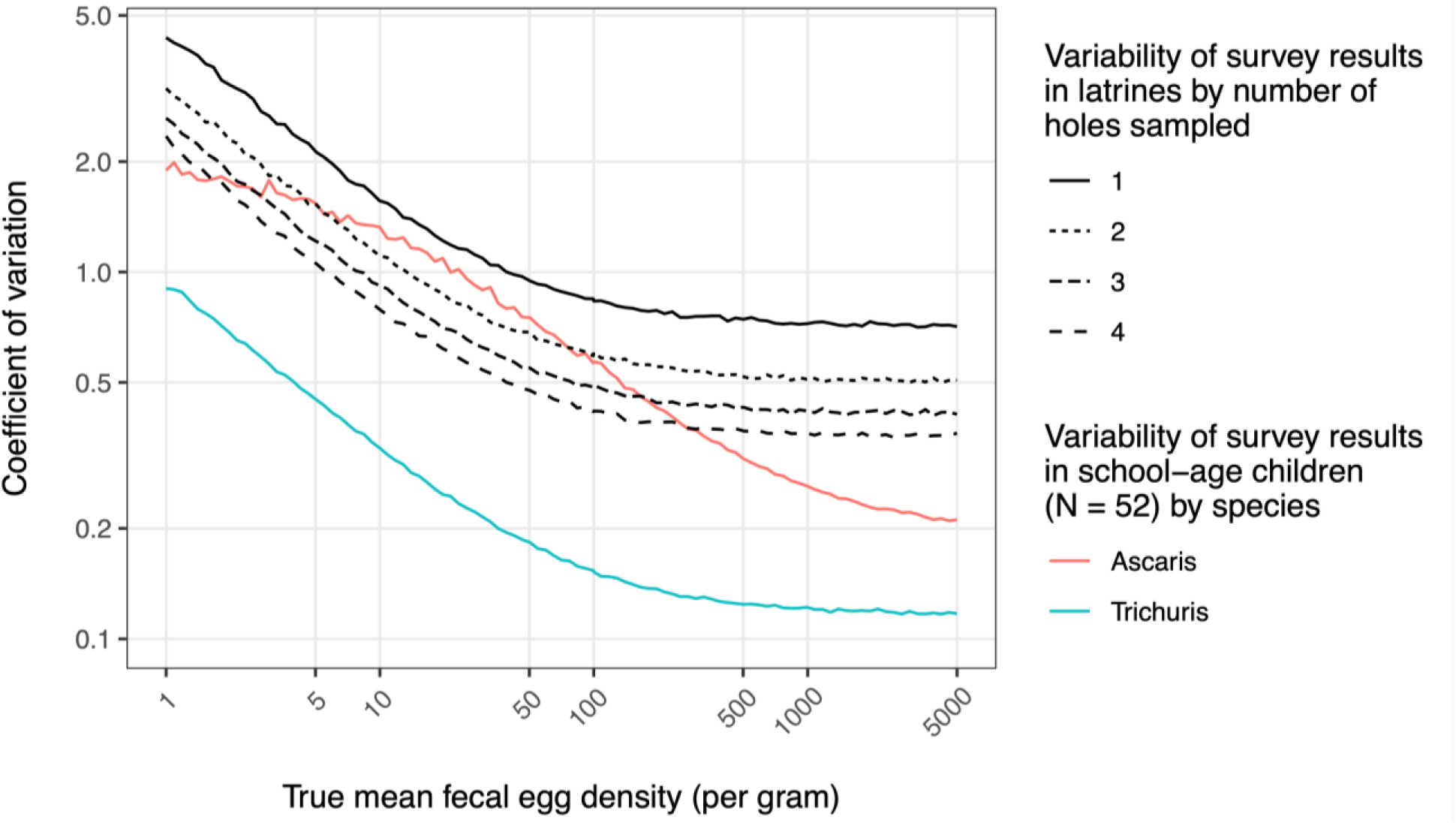
Association between average and variance of egg counts across slides, processed samples, and pit latrine holes in field settings. Variability of survey results is expressed in terms of the coefficient of variation of the school-level mean egg count across repeated survey results (y-axis), based on 10,000 repeated simulations. For lack of prior data, we assumed that the true egg density in sludge from latrines is the same as the true average egg density in faeces from children (x-axis). Survey results in school-age children are based on duplicate Kato-Katz, using a single well-homogenised stool sample per child, adopting a previously developed simulation framework for soil-transmitted helminths in school-age children [35]. Compared to Kato-Katz applied directly to children’s stools, sludge testing with the Fluke Catcher method was assumed to recover only 50% of the eggs (based on the spiking experiments; Fig 4).

First, the CV changes with endemicity. The variability is highest when the mean egg density is at its lowest (5 EPG), then steeply drops as a function of increasing true mean egg density, after which it stabilizes from 50 EPG onwards. Second, when aiming to reduce the variability in survey results it is better to sample more squat holes than to repeatedly process the same fecal sludge sample. For instance, assuming a true mean egg intensity of 100 EPG, the CV was 0.83 when surveys are based on a single slide from a single fecal sludge sample and squat hole. Increasing the number of sampled squat holes to four (still single sample prep and one slide per sample prep) reduced the CV to 0.42, whereas increasing the number of sample preps to four (still single slide per sample prep) only reduced the CV to 0.63. These findings are directly explained by the finding that variability in egg counts in the field study (previous section) were mostly explained by variability between squat holes (i.e., highest CV, **Table 2**).

Generally, the results of stool-based surveys among children (red and blue lines in **Fig 7**) were less variable compared to latrine-based surveys. However, the CV of latrine-based surveys did not exceed that of stool-based surveys for *Ascaris* when the true mean fecal density was below 100 EPG and at least 3 to 4 squat holes were sampled. For detection of *Trichuris* eggs, surveys among children yielded consistently more precise results (lower CVs) than latrine-based surveys.

## Discussion

### Fecal sludge sampling devices can be locally made

Designing and field testing of prototypes for fecal sludge sampling from school pit latrines showed that, simple devices can be constructed from readily and locally available materials and adapted to accommodate variable pit latrine structures. The earlier prototype devices we designed faced several operational challenges that required ad-hoc modifications, including narrower fecal sludge sampling containers and extended sampling poles, which remained feasible under field conditions. The modified fecal sludge sampling prototypes were generally successful, except for a few pit latrines with dried/solid sludge types and insufficient sludge volume (e.g., newly constructed, recently emptied and infrequently used school pit latrines). These limitations indicate that latrine-based survey might not be universally applicable across all sanitation contexts.

### Our modified Fluke Catcher method is simple, cheap and has moderate performance

To develop a streamlined fecal sludge sample processing, we reviewed existing protocols for the detection and quantification of STH eggs from sludge samples. Although several methods have been described [30,37,42,45,46], many rely on labor-intensive procedures, expensive equipment, and/or reagents/solutions which are often expensive or toxic for both humans and environment. In the present study, we combined the Fluke Catcher with a 20 µm pore size sieve for the detection and quantification of STH eggs in fecal sludge, and by doing so omitting the need of chemical solutions [25,30]. The cost of the modified Fluke Catcher is estimated to be 478.50 EUR, which is 94% less expensive than that of our previously described egg-count method to detect and quantify STH in soil (8,000.00 EUR) [25]. The evaluation of the modified Fluke Catcher method showed that the analytical performance ranged from 56 to 85%, the performance being highest when four slides were examined and when eggs were spiked in liquid sludge samples and being lowest when one slide was examined and when eggs were spiked in semi-solid samples. The egg recovery rate depended on the consistency of the fecal sludge consistency, with more eggs being recovered in liquid fecal sludge (85%; *vs.* slurry (65%) *vs.* semi-solid sludge (56%)). These differences across fecal sludge consistency can be explained by the ‘sticky’ nature of *Ascaris eggs*, which can easily get trapped in denser fecal sludge and therefore get lost in semi-solid vs. more liquid samples [47,48]. A review study on recent advances in quantification of STH eggs from environmental samples also reported similar egg recovery rates (57 – 80%) [24], indicating that our egg counting method has a moderate performance. Of course, we only conducted experiments with *Ascaris* eggs (which was the most abundant STH in the area), and thus extrapolation on the analytical performance of our modified Fluke Catcher method to other STH and helminthiases should be done with care. In addition, the method has some important limitations. The need for a centrifuge is one of them. Exploring alternatives such as passive sedimentation would therefore be recommended [24]. A potential tool that could assist in this is the FECPAK^G2^ sedimenter, which has already been optimized for stool by Ayana *et al*. [31]. Similarly, our fecal sludge sample processing protocol is an open system, meaning that further methodological modifications are required to transform it into a closed system to minimize biosafety risks [49].

### Variation in school abundance underscores the potential of a latrine-based survey

We observed six medically important helminths (*Ascaris, Trichuris*, *S. mansoni*, *E. vermicularis*, *H. nana* and *Taenia* spp.). We also detected hookworm-like eggs, but in absence of any molecular analysis, we did not feel comfortable in drawing conclusions on the presence of this STH. Noteworthy is the observed variation in abundance (mean egg counts per gram of fecal sludge total solid) across schools for *Ascaris*, *Trichuris* and *S. mansoni*). Similarly, historical data on a stool-based survey across 10 schools in the same study area (2015 and 2018; [32]) indicated a wide range in school prevalence and intensity for each of the STHs (see **Table S2**). **Fig S3** plots the historical prevalence data with the abundance in sludge samples for six schools involved in the present study, underscoring the potential of a latrine-based survey. In a follow-up study we will conduct a head-to-head comparison of a latrine-based and a stool-based (individual stool samples collected from children) survey across 25 schools (52 children per school) in Jimma Zone (Ethiopia).

### Insights into sources of variation essential to determine sampling and analysis strategy

To determine the strategy to sample (required number of squat holes per latrine, depth of sample collection) and analysis fecal sludge samples (the number sample preps and slides) for this follow-up study, in the current study, we prioritized understanding the sources of variation in helminth egg counts. The main source of egg count variations was due to difference in egg counts between squat holes, followed by variation in sample preps and microscopists. This pattern of better increasing sampling efforts over analysis efforts is in line with the study by Koottatep *et al*. [41]. This study reported that in a fecal sludge treatment plant multiple sampling at different points provides more representative estimates than a single point sampling. In contrast, no significant differences in abundance were observed between boys’ and girls’ pit latrines and depth of sampling. The subsequent simulation study indicated that a latrine-based survey was only as precise as a stool-based survey for *Ascaris* and when the fecal egg density was below 100 EPG. For the detection of *Trichuris* eggs, surveys among children yielded consistently more precise results (lower CVs) than latrine-based surveys. This difference may reflect that the CV for *Trichuris* egg counts might be expected to be lower than what currently shown here, as our estimates of variability of latrine-based surveys are mostly driven by *Ascaris*.

## Conclusion

This study demonstrates the design and feasibility of low cost locally manufactured devices for fecal sludge sampling in school pit latrines and provides a detailed evaluation of the analytical performance of an egg-counting method under laboratory conditions. It also identified the most important sources of variation in egg counts, allowing to define a more evidence-based sampling collection and analysis strategy. Together, the findings support the potential of latrine-based survey as an alternative to stool-based surveys, while also highlighting important methodological constraints that must be addressed to ensure reliable estimation of STH egg concentrations. Finally, we suggested sampling two to three holes per pit latrine from top layer of sludge, one sample processing and one slide (representing a volume of 50 µL) examination to validate our diagnostic method in 25 primary schools in Jimma Zone, Southwest Ethiopia.

## Acknowledgements

This work was supported, in whole or in part, by the Gates Foundation [Investment ID 049001]. The conclusions and opinions expressed in this work are those of the author(s) alone and shall not be attributed to the Foundation. Under the grant conditions of the Foundation, a Creative Commons Attribution 4.0 License has already been assigned to the Author Accepted Manuscript version that might arise from this submission. Please note works submitted as a preprint have not undergone a peer review process.

We thank Wim Roose for his valuable contributions to the design and improvement of the fecal sludge sampling prototypes.

## References

1. Melvin DM, Brooke MM, Sadun EH. Common Intestinal Helminths of Man: Life Cycle Charts. US Department of Health, Education, and Welfare, Public Health Service, Communicable Disease Center; 1965. CDC Information Center, Atlanta, GA 30333. 3–8 p. Available from: https://stacks.cdc.gov/view/cdc/7658/cdc_7658_DS1.pdf.

2. World Health Organization. Eliminating soil-transmitted helminthiases as a public health problem in children. Progress report 2001 – 2010 and strategic plan 2011−2020. WHO, Geneva, 2012. Available from: https://www.who.int/publications/i/item/9789241503129.

3. Prüss-Ustün A, Wolf J, Bartram J, Clasen T, Cumming O, Freeman MC, et al. Burden of disease from inadequate water, sanitation and hygiene for selected adverse health outcomes: An updated analysis with a focus on low- and middle-income countries. Int J Hyg Environ Health. 2019 Jun 1;222(5):765–77.

4. Blouin B, Casapia M, Joseph L, Gyorkos TW. A longitudinal cohort study of soil-transmitted helminth infections during the second year of life and associations with reduced long-term cognitive and verbal abilities. PLoS Negl Trop Dis. 2018;12(7): e0006688.

5. Hotez PJ, Bundy DAP, Beegle K, Brooker S, Drake L, Silva N De, et al. Helminth Infections: Soil-transmitted Helminth Infections and Schistosomiasis. In: Jamison DT, Breman JG, Measham AR, et al., editors. Disease Control Priorities in Developing Countries. 2nd edition. Washington (DC): The International Bank for Reconstruction and Development / The World Bank; 2006. Chapter 24. Available from: https://www.ncbi.nlm.nih.gov/books/NBK11748/. Co-published by Oxford University Press, New York.

6. Aderoba AK, Iribhogbe OI, Olagbuji BN, Olokor OE, Ojide CK, Ande AB. Prevalence of helminth infestation during pregnancy and its association with maternal anemia and low birth weight. International Journal of Gynecology & Obstetrics. 2015 Jun 1;129(3):199–202.

7. World Health Organization. Guideline: preventive chemotherapy to control soil-transmitted helminth infections in at-risk population groups. Geneva: WHO; 2017. Licence: CC BY-NC-SA 3.0 IGO. Available from: https://wkc.who.int/resources/publications/i/item/9789241550116.

8. Anderson R, Truscott J, Hollingsworth TD. The coverage and frequency of mass drug administration required to eliminate persistent transmission of soil-transmitted helminths. Philosophical Transactions of the Royal Society B: Biological Sciences. 2014 Jun 19;369(1645):20130435.

9. Montresor A, Mwinzi P, Mupfasoni D, Garba A. Reduction in DALYs lost due to soil-transmitted helminthiases and schistosomiasis from 2000 to 2019 is parallel to the increase in coverage of the global control programmes. PLoS Negl Trop Dis. 2022;16(7): e0010575.

10. World Health Organization. 2030 targets for soil-transmitted helminthiases control programmes. Geneva: WHO; 2019. Licence: CC BY-NC-SA 3.0 IGO. Available from: https://www.who.int/publications/i/item/9789240000315.

11. School Health Integrated Programming. Guidelines for School-based Deworming Programs: Information for policy-makers and planners on conducting deworming as part of an integrated school health program. Partnership for Child Development, Imperial College, London 2016. Available from: https://resourcecentre.savethechildren.net/document/guidelines-school-based-deworming-programs-information-policy-makers-and-planners-conducting.

12. Giardina F, Coffeng LE, Farrell SH, Vegvari C, Werkman M, Truscott JE, et al. Sampling strategies for monitoring and evaluation of morbidity targets for soil-transmitted helminths. PLoS Negl Trop Dis. 2019; 13(6): e0007514.

13. Sturrock HJW, Gething PW, Ashton RA, Kolaczinski JH, Kabatereine NB, Brooker S. Planning schistosomiasis control: Investigation of alternative sampling strategies for Schistosoma mansoni to target mass drug administration of praziquantel in East Africa. Int Health. 2011;3(3):165–175.

14. Kura K, Truscott JE, Collyer BS, Phillips A, Garba A, Anderson RM. The observed relationship between the degree of parasite aggregation and the prevalence of infection within human host populations for soil-transmitted helminth and schistosome infections. Trans R Soc Trop Med Hyg. 2022;116(12):1226–1229.

15. Coffeng LE, Malizia V, Vegvari C, Cools P, Halliday KE, Levecke B, et al. Impact of Different Sampling Schemes for Decision Making in Soil-Transmitted Helminthiasis Control Programs. J Infect Dis. 2020 Jun 11;221(Suppl 5):S531–S538.

16. Speich B, Knopp S, Mohammed KA, Khamis S, Rinaldi L, Cringoli G, et al. Comparative cost assessment of the Kato-Katz and FLOTAC techniques for soil-transmitted helminth diagnosis in epidemiological surveys. Parasit Vectors. 2010 Aug 14;3:71.

17. Leta GT, French M, Dorny P, Vercruysse J, Levecke B. Comparison of individual and pooled diagnostic examination strategies during the national mapping of soil-transmitted helminths and Schistosoma mansoni in Ethiopia. PLoS Negl Trop Dis. 2018 Sep 1; 12(9): e0006723.

18. World Health Organization. Assessing schistosomiasis and soil-transmitted helminthiases control programmes: monitoring and evaluation framework. Geneva: WHO; 2024. Licence: CC BY-NC-SA 3.0 IGO. Available from: https://www.who.int/publications/b/73248.

19. Minnery M, Okoyo C, Morgan G, Wang A, Johnson O, Fronterre C, et al. Cost-effectiveness of comparative survey designs for helminth control programs: Posthoc cost analysis and modelling of the Kenyan national school-based deworming program. PLoS Negl Trop Dis. 2024 Dec 1;18(12): e0011583.

20. Mekonnen Z, Meka S, Ayana M, Bogers J, Vercruysse J, Levecke B. Comparison of Individual and Pooled Stool Samples for the Assessment of Soil-Transmitted Helminth Infection Intensity and Drug Efficacy. PLoS Negl Trop Dis. 2013; 7(5): e2189.

21. World Health Organization. Diagnostic target product profiles for monitoring and evaluation of soil-transmitted helminth control programs. Geneva: WHO; 2021. Licence: CC BY-NC-SA 3.0 IGO. Available from: https://www.who.int/publications/i/item/9789240031227.

22. Santarpia JL, Klug E, Ravnholdt A, Kinahan SM. Environmental sampling for disease surveillance: Recent advances and recommendations for best practice. J Air Waste Manag Assoc. 2023 Jun;73(6):434–461.

23. Colenutt C, Brown E, Nelson N, Wadsworth J, Maud J, Adhikari B, et al. Environmental Sampling as a Low-Technology Method for Surveillance of Foot-and-Mouth Disease Virus in an Area of Endemicity. Appl Environ Microbiol. 2018 Aug 1;84(16):e00686–18.

24. Amoah ID, Singh G, Stenström TA, Reddy P. Detection and quantification of soil-transmitted helminths in environmental samples: A review of current state-of-the-art and future perspectives. Acta Trop. 2017 May;169:187–201.

25. Tadege B, Mekonnen Z, Dana D, Sharew B, Dereje E, Loha E, et al. Assessment of environmental contamination with soil-transmitted helminths life stages at school compounds, households and open markets in Jimma Town, Ethiopia. PLoS Negl Trop Dis. 2022 Apr 4;16(4):e0010307.

26. Manuel M, Amato HK, Pilotte N, Chieng B, Araka SB, Siko JEE, et al. Soil surveillance for monitoring soil-transmitted helminths: Method development and field testing in three countries. PLoS Negl Trop Dis. 2024 Sep 6;18(9):e0012416.

27. Siko JEE, Dahmer KJ, Manoharan ZZ, Muthukumar A, Amato HK, LeBoa C, et al. Environmental surveillance of soil-transmitted helminths and other enteric pathogens in settings without networked wastewater infrastructure. PLOS Water. 2025;4(1):e0000337.

28. Provinos. Sheep and other animals: Provinos Botvanger – simple manure testing for liver fluke. Online store Protanso & Provinos. Netherlands, 2026. Available from: https://www.provinos.nl/.

29. Retsch®. Product – Test Sieves: woven wire mesh sieves. Germany, 2026. Available from: https://www.retsch.com/.

30. Velkushanova K. Strande L. Ronteltap M. Koottatep T. Brdjanovic D. Buckley C. (eds.). Methods for Faecal Sludge Analysis. IWA Publishing, London, UK; 2021. ISBN:9781780409115.

31. Ayana M, Vlaminck J, Cools P, Ame S, Albonico M, Dana D, et al. Modification and optimization of the FECPAKG2 protocol for the detection and quantification of soil-transmitted helminth eggs in human stool. PLoS Negl Trop Dis. 2018 Oct 15;12(10):e0006655.

32. Dana D, Vlaminck J, Ayana M, Tadege B, Mekonnen Z, Geldhof P, et al. Evaluation of copromicroscopy and serology to measure the exposure to Ascaris infections across age groups and to assess the impact of 3 years of biannual mass drug administration in Jimma Town, Ethiopia. PLoS Negl Trop Dis. 2020 Apr 13;14(4):e0008037.

33. Debela TH, Beyene A, Tesfahun E, Getaneh A, Gize A, Mekonnen Z. Fecal contamination of soil and water in sub-Saharan Africa cities: The case of Addis Ababa, Ethiopia. Ecohydrology and Hydrobiology. 2018 Apr 1;18(2):225–230.

34. Coffeng LE, Graham M, Browning R, Kura K, Diggle PJ, Denwood M, et al. Improving the Cost-efficiency of Preventive Chemotherapy: Impact of New Diagnostics on Stopping Decisions for Control of Schistosomiasis. Clin Infect Dis. 2024 Apr 25;78(Suppl 2):S153–S159.

35. Kazienga A, Levecke B, Leta GT, de Vlas SJ, Coffeng LE. general framework to support cost-efficient survey design choices for the control of soil-transmitted helminths when deploying Kato-Katz thick smear. PLoS Negl Trop Dis. 2023 Jun 22;17(6):e0011160.

36. Gates Foundation. Sampling for faecal sludge and other liquid wastes in emergency settings [version 1; not peer reviewed]. Gates Open Res 2021; 5:27 (document). Available from: 10.21955/gatesopenres.1116730.1.

37. Nabateesa S, Zziwa A, Kabenge ISA, Kambugu R, Wanyama J, Komakech AJ. Occurrence and survival of pathogens at different sludge depths in unlined pit latrines in kampala slums. Water SA. 2017 Oct 1;43(4):638–645.

38. Capone D, Chigwechokha P, de los Reyes FL, Holm RH, Risk BB, Tilley E, et al. Impact of sampling depth on pathogen detection in pit latrines. PLoS Negl Trop Dis. 2021 Mar 2;15(3):e0009176.

39. Capone D, Berendes D, Cumming O, Knee J, Nalá R, Risk BB, et al. Analysis of Fecal Sludges Reveals Common Enteric Pathogens in Urban Maputo, Mozambique. Environ Sci Technol Lett. 2020;7(12):889–895.

40. Nzouebet WAL, Noumsi IMK, Rechenburg A. Prevalence and diversity of intestinal helminth eggs in pit latrine sludge of a tropical urban area. Journal of Water Sanitation and Hygiene for Development. 2016;6(4):622–630.

41. Koottatep T, Ferré A, Chapagain S, Fakkaew K, Strande L. Faecal sludge sample collection and handling. In: Velkushanova K, Strande L, Ronteltap M, Koottatep T, Brdjanovic D, Buckley C, editors. Methods for Faecal Sludge Analysis. London, UK: IWA Publishing; 2021. p. 55–84.

42. U.S. Environmental Protection Agency (EPA). Environmental Regulations and Technology: Control of Pathogens and Vector Attraction in Sewage Sludge, (Including Domestic Septage) Under 40 CFR Part 503. Cincinnati, OH: U.S. EPA, Office of Research and Development, National Risk Management Research Laboratory, Center for Environmental Research Information. Revised July 2003. EPA/625/R-92/013. Available from: https://www.epa.gov/sites/default/files/2015-04/documents/control_of_pathogens_and_vector_attraction_in_sewage_sludge_july_2003.pdf.

43. Rahman M, Islam M, Doza S, Naser AM, Shoab AK, Rosenbaum J, et al. Higher helminth ova counts and incomplete decomposition in sand-enveloped latrine pits in a coastal sub-district of Bangladesh. PLoS Negl Trop Dis. 2022 Jun 23;16(6):e0010495.

44. Kumwenda S, Msefula C, Kadewa W, Diness Y, Kato C, Morse T, et al. Is there a difference in prevalence of helminths between households using ecological sanitation and those using traditional pit latrines? A latrine based cross sectional comparative study in Malawi. BMC Res Notes. 2017 Jun 9;10(1):200.

45. Moodley P, Archer A, Hawksworth D. Standard Methods for the Recovery and Enumeration of Helminth Ova in Wastewater, Sludge, Compost and Urine-Diversion Waste in South Africa. Water Research Commission, Pretoria, South Africa, 2008. WRC Report No. TT322/08. Available from: https://www.wrc.org.za/wp-content/uploads/mdocs/TT%20322-web.pdf.

46. Bowman DD, Little MD, Reimers RS. Precision and accuracy of an assay for detecting Ascaris eggs in various biosolid matrices. Water Res. 2003 May;37(9):2063–72.

47. Jiménez B, Maya C, Galván M. Helminth ova control in wastewater and sludge for advanced and conventional sanitation. Water Sci Technol. 2007;56(5):43–51.

48. Naidoo D, Archer CE. *Ascaris suum* Egg Recovery from Sludge Samples after Phase Extraction. J Parasitol. 2024 Jul 1;110(4):295–299.

49. World Bank, ILO, WaterAid, and WHO. Health, Safety and Dignity of Sanitation Workers: An Initial Assessment. World Bank, Washington, DC, USA; 2019. Available from: https://documents1.worldbank.org/curated/en/316451573511660715/pdf/Health-Safety-and-Dignity-of-Sanitation-Workers-An-Initial-Assessment.pdf.

50. Stuyver LJ, Levecke B. The role of diagnostic technologies to measure progress toward WHO 2030 targets for soil-transmitted helminth control programs. PLoS Negl Trop Dis. 2021 Jun 3;15(6):e0009422.

51. Urdea M, Penny LA, Olmsted SS, Giovanni MY, Kaspar P, Shepherd A, et al. Requirements for high impact diagnostics in the developing world. Nature. 2006 Nov 23;444 (Suppl 1):73–9.

